# Heritability and genome-wide association study of diffusing capacity of the lung

**DOI:** 10.1101/277343

**Authors:** Natalie Terzikhan, Fangui Sun, Fien M. Verhamme, Hieab H.H. Adams, Daan Loth, Ken R. Bracke, Bruno H. C. Stricker, Lies Lahousse, Josée Dupuis, Guy G. Brusselle, George T. O’Connor

## Abstract

**Background:** Although several genome wide association studies (GWAS) have investigated the genetics of pulmonary ventilatory function, little is known about the genetic factors that influence gas exchange.

**Aim:** To investigate the heritability of, and genetic variants associated with the diffusing capacity of the lung.

**Methods:** GWAS was performed on diffusing capacity, measured by carbon monoxide uptake (DLCO) and per alveolar volume (DLCO/VA) using the single-breath technique, in 8,372 individuals from two population-based cohort studies, the Rotterdam Study and the Framingham Heart Study. Heritability was estimated in related (n=6,246) and unrelated (n=3,286) individuals.

**Results:** Heritability of DLCO and DLCO/VA ranged between 23% and 28% in unrelated individuals and between 45% and 49% in related individuals. Meta-analysis identified a genetic variant in *GPR126* that is significantly associated with DLCO/VA. Gene expression analysis of *GPR126* in human lung tissue revealed a decreased expression in patients with COPD and subjects with decreased DLCO/VA.

**Conclusion:** DLCO and DLCO/VA are heritable traits, with a considerable proportion of variance explained by genetics. A functional variant in *GPR126* gene region was significantly associated with DLCO/VA. Pulmonary *GPR126* expression was decreased in patients with COPD.

## Introduction

The respiratory system can be separated functionally into two zones. The first one is the conducting zone, which includes the trachea, bronchi, bronchioles, and terminal bronchioles and which is functional in ventilation, i.e. conducting the air in and out of the lungs. The second zone is the respiratory zone, which consists of the respiratory bronchioles, alveolar ducts and alveoli, the site where oxygen and carbon-dioxide are exchanged between the lungs and the blood.

Different pulmonary function tests are available that measure these distinct functions of ventilation and gas exchange. These tests help to evaluate and manage patients with respiratory symptoms and diseases, and include spirometry, measurements of lung volumes, and the diffusing capacity of the lung for carbon monoxide (DLCO). The latter, also known as transfer factor of the lung for CO, provides a quantitative measure of gas transfer in the lung (1, 2) and reflects processes in the alveolar compartment and pulmonary microcirculation.

The DLCO provides clinical insights complimentary to those obtained by spirometry and lung volume measurements, for example, in discriminating asthma from chronic obstructive pulmonary disease (COPD), to identify causes of hypoxemia or dyspnoea, and to monitor patients with interstitial lung disease (3). DLCO is decreased in patients with emphysema due to a decrease in the total surface area of the lung and the loss of capillary beds (1, 4). In contrast to the abundance of genome-wide association studies (GWAS) investigating genetic variation of spirometry measures (5–8), the heritability of, and genetic influences on DLCO, are largely unknown.

Therefore, we first investigated the heritability of DLCO to understand which proportion of the variance in DLCO can be explained by genetics. Next, we performed a GWAS, to identify genetic variants affecting the variability in DLCO, using data from two prospective population-based cohorts; the Rotterdam Study and the Framingham Heart Study. Finally, we investigated the expression of the lead GWAS association in lung tissue of individuals with COPD and (non-smoking and smoking) controls.

## Methods

In this section the methods will be described briefly. Please see **Supplemental Methods** in the Online Data Supplement for more detailed information.

### Setting

The present meta-analysis combined results from two population-based studies, i.e. the Rotterdam Study and the Framingham Heart Study. The Rotterdam Study (9) is an ongoing prospective population-based cohort study that includes 3 cohorts encompassing 14,926 participants aged ≥ 45 years, living in the Netherlands. DLCO was measured between 2009-2013.

The Framingham heart study is a population-based family study that recruited residents of Framingham, Massachusetts starting from 1948. DLCO was measured at the 8th and 9th examinations of the Offspring Cohort (2005-2008 and 2011-2014) and the 1st and 2nd examinations of the Third Generation Cohort (2002-2005 and 2008-2011). For participants with measurements at both time points, we analyzed the later measurement.

### Lung function

DLCO (mmol/min/kPA) and alveolar volume (VA) were measured by the single breath technique in accordance with ERS / ATS guidelines (2). The DLCO per alveolar volume (DLCO/VA; mmol/min/kPA/liter) was calculated by dividing the DLCO by VA. Analyses were restricted to participants with two interpretable and reproducible measurements of DLCO and DLCO/VA.

### Heritability analysis

Heritability was defined as the ratio of trait variance due to additive genetic effects to the total phenotypic variance after accounting for covariates. In the Rotterdam Study GCTA software (10) was used to estimated heritability in unrelated individuals. In the Framingham Heart Study SOLAR software (11) was used to estimate heritability based on familial relationships. Analyses were adjusted for age, sex and principal components of genetic relatedness ((PC), in GCTA only). Additional adjustment for current and former smoking were done in a subsequent analysis.

### GWAS analyses

A GWAS was performed for both phenotypes DLCO and DLCO/VA using ProbABEL (version 0.4.4). Variants with imputation quality (R^2^) < 0.3 and minor allele frequency (MAF) < 0.01 were excluded from the analyses. Linear regression was conducted for each SNP, assuming additive model. All analyses were adjusted for age, sex and PC (Rotterdam Study only) in model 1 and additionally adjusted for smoking, weight and height in model 2. A random effect was added to the model to account for familial relationship in the Framingham Heart Study analyses. Data were meta-analyzed using METAL software and were adjusted for genomic control. Genome-wide significance threshold was set at P-value < 5 x10^−8^ and for suggestive associations at P-value 5 × 10^−7^. Quantile-quantile plots, Manhattan plots and regional plots were generated using the R software. Analyses were repeated after 1) correction for haemoglobin in the Rotterdam Study and 2) additional adjustment for FEV_1_/FVC.

### Follow-up analyses

Several steps were taken in order to explore the functionality of the variants and genes of interest, and to associate those newly identified loci to clinically relevant disease outcomes. 1) Genetic correlations were investigated, 2) Genetic overlap was investigated with SNPs that are significantly related to COPD (12) and emphysema (13). 3) Posterior probability of causality of the lead SNP was calculated using FINEMAP software (14) .4) The regulatory function of the lead SNP was explored on the Haploreg server. 5) The effect of the lead SNP on mRNA expression was checked (Expression Quantitative Trait Loci (eQTL) analysis), using lung tissue dataset from Genotype Tissue Expression (GTEx) portal (see **URLs**). 6) Tissue-specific gene expression was checked in GTEx portal and 7) Finally, mRNA expression of the *GPR126* gene was analysed in lung tissue (using real-time PCR) of 92 patients with or without COPD.

## Results

### The study cohorts and participant characteristics

The general characteristics of the study populations (the Rotterdam Study and the Framingham Heart Study) are shown in **Table 1**. The mean age was 67.3 (SD 8.0 yr) years in the Rotterdam Study and 52.8 (SD 14.8 yr) years in the Framingham Heart Study. **Figure 1** shows the study flow of participants that were included in this study.

**Table 1:** General characteristics of the study populations

|  | RS | FHS |
| --- | --- | --- |
|  | N= 2,574 | N= 5,798 |
| Age, years | 67.3 (8.0) | 52.7 (14.8) |
| Female, % | 51.9 | 53.9 |
| Weight, Kg | 80.5 (14.9) | 79.7 (18.5) |
| Height, cm | 170.6 (9.2) | 168.9 (9.5) |
| Former smokers, % | 55.4 | 39.9 |
| Current smokers, % | 11.5 | 10.9 |
| Never smokers, % | 33.1 | 49.2 |
| DLCO (mmol/min/kPA) | 8.0 (1.8) | 8.3 (2.3) |
| DLCO corrected for Hb (mmol/min/kPA) | 7.9 (1.7) | NA |
| DLCO/VA (mmol/min/kPA/VA) | 1.5 (0.2) | 1.5 (0.2) |
| DLCO/VA corrected for Hb (mmol/min/kPA/VA) | 1.5 (0.2) | NA |
| FEV<sub>1</sub> (L) | 2.8 (0.7) | 3.1 (0.9) |
| FEV<sub>1</sub>, % predicted | 105.3 (19.8) | 99.2 (14.9) |
| FVC (L) | 3.7 (1.0) | 4.2 (1.1) |
| FVC, % predicted | 110.1 (17.6) | 102.7 (13.5) |
| FEV<sub>1</sub>/FVC, % | 76.4 (7.1) | 75.4 (6.9) |

DLCO: Diffusing capacity of the lung for carbon monoxide; DLCO/VA: Diffusing capacity of the lung for carbon monoxide by alveolar volume; FEV<sub>1</sub>: Forced expiratory volume during the first second; FHS: Framingham Heart Study; FVC: Forced vital capacity; Hb: Haemoglobin; RS: Rotterdam Study
Values are means (standard deviation) for continuous variables or percentages for dichotomous variables.

**Figure. 1:**
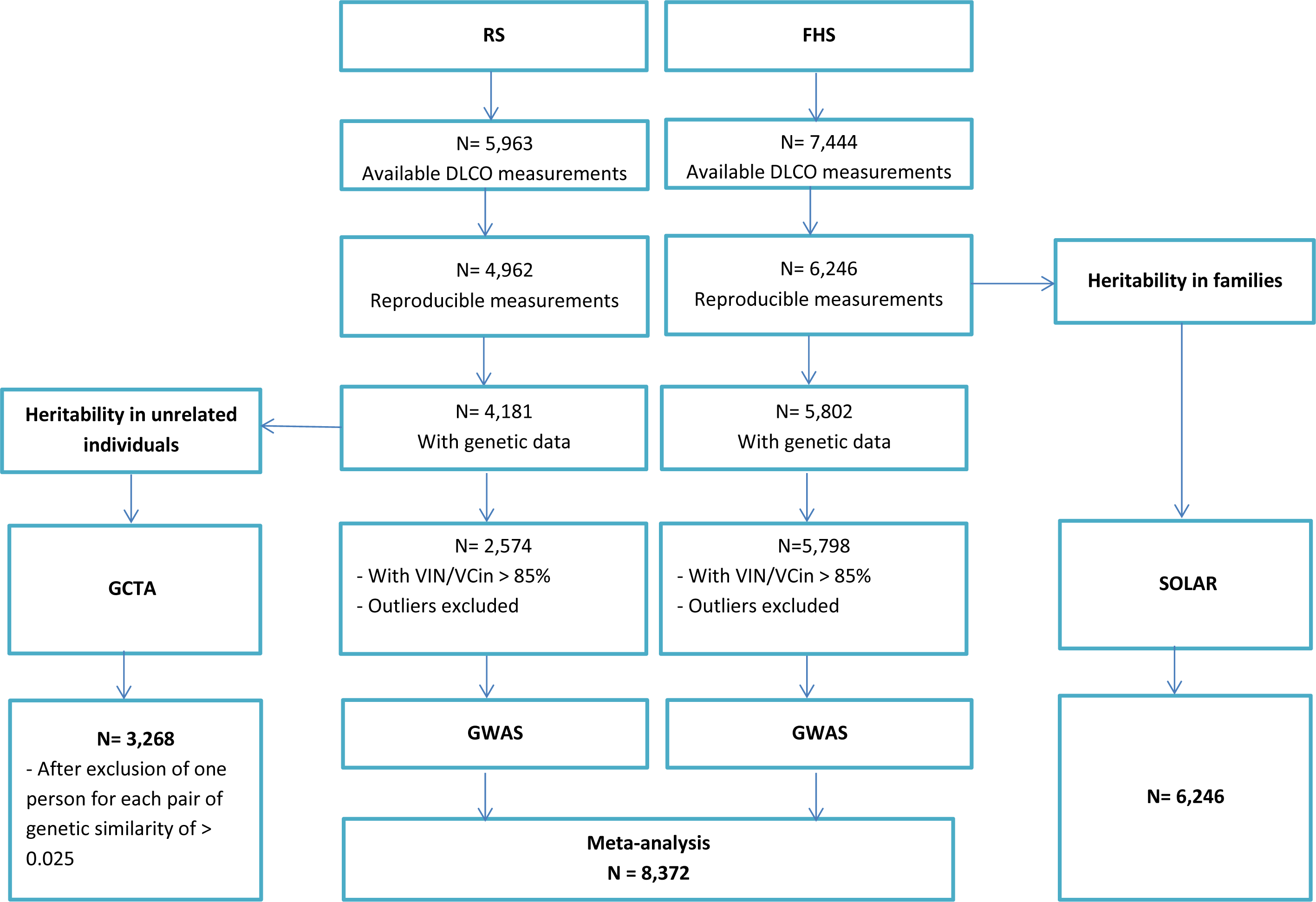
Flowchart of study participants. DLCO: Diffusing capacity of carbon monoxide; FHS; the Framingham Heart Study; GWAS; genome wide association study; GCTA: Genome-wide Complex Trait Analysis software; N= number of participants; RS: the Rotterdam Study; SOLAR: Sequential Oligogenic Linkage Analysis Routines package; VA: Alveolar volume; VCin: vital capacity measured during maximal inspiration; VIN: inspiratory volume.

### Heritability

Heritability was estimated in two ways, the first one was by using the Rotterdam Study data with unrelated individuals, with a total number of 3,286 participants with genetic data and interpretable measurements of DLCO. The second was by using data from the Framingham Heart Study to estimate heritability based on familial relationships in 6,246 participants with interpretable measurements of DLCO (**Figure 1**). In **Table 2** heritability estimates for DLCO and DLCO/VA are presented. In unrelated individuals, we found an age- and sex- and PC-adjusted heritability for DLCO of 23%, and a heritability of 28% after additional adjustment for current and past smoking. Similar heritability estimates were found for DLCO/VA with 24% after adjustment for age, sex and PC, and 25% after additional adjustment for smoking. In the Framingham Heart Study, investigating individuals with known familial relationships, we found an age- and sex-adjusted heritability for DLCO of 49%, and a heritability of 47% after additional adjustment for current and past smoking. Heritability estimates for DLCO/VA were 45% after adjustment for age, sex, and 46% after additional adjustment for current and past smoking.

**Table 2:** Heritability of diffusing capacity of the lung

| Model* | Rotterdam Study (RS) |  |  |  | Framingham Heart Study (FHS) |  |  |  |
| --- | --- | --- | --- | --- | --- | --- | --- | --- |
|  | *N=3,286 unrelated individuals* |  |  |  | *N=6,246 with known familial relationships* |  |  |  |
|  | DLCO |  | DLCO/VA |  | DLCO |  | DLCO/VA |  |
|  | **h<sup>2</sup> (SE)** | **P-value** | **h<sup>2</sup> (SE)** | **P-value** | **h<sup>2</sup> (SE)** | **P-value** | **h<sup>2</sup> (SE)** | **P-value** |
| **1** | 0.23 (0.10) | 0.01 | 0.24 (0.10) | 0.009 | 0.49 (0.03) | $2.3 \times 10^{-106}$ | 0.45 (0.03) | $5.0 \times 10^{-82}$ |
| **2** | 0.28 (0.10) | 0.002 | 0.25 (0.10) | 0.0075 | 0.47 (0.03) | $8.5 \times 10^{-100}$ | 0.46 (0.03) | $7.6 \times 10^{-84}$ |

DLCO: Diffusing capacity of the lung for carbon monoxide; DLCO/VA: Diffusing capacity of the lung for carbon monoxide by alveolar volume; FHS: The Framingham Heart Study; h<sup>2</sup>: heritability estimate; RS: The Rotterdam Study; SE: Standard error.
\* Model 1: adjusted for age, sex and principal components of genetic relatedness (RS only); Model 2: adjusted for age, sex, smoking and principal components of genetic relatedness (RS only).

### Genetic variants associated with diffusing capacity

We performed GWAS on DLCO and DLCO/VA in the Rotterdam Study (n=2,574) and the Framingham Heart Study (n= 5,798), and subsequently meta-analysed both cohorts (n= 8,372). All variants with a p-value below 5 × 10^−6^ at the meta-analysis stage are presented in **Table 3**. The corresponding quantile-quantile plots are presented in **Figure E1** in the Online Data Supplement. GWAS results of the separate cohorts with (P-value < 5 × 10^−6^), are presented in **Tables E1 and E2** in the Online Data Supplement.

**Table 3:** Independent genetic variants that are significantly or suggestively associated with DLCO or DLCO/VA at meta-analysis level.

| Trait | SNP | Chr:Pos | Gene ‡ | A1/A2 | RS (n=2,574) |  | FHS (n=5,798) |  | RS and FHS (n=8,376) |  |
| --- | --- | --- | --- | --- | --- | --- | --- | --- | --- | --- |
|  |  |  |  |  | B | P-value | B | P-value | B | P-value |
| **DLCO\*** | - | - | - | - | - | - | - | - | - | - |
| **DLCO†** | rs1665630 | 10:73426862 | CDH23 | T/C | .11 | 6.4x10<sup>-4</sup> | .10 | 9.1x10<sup>-5</sup> | -.02 | 2.8x10<sup>-7</sup> |
|  | rs2423124 | 20:5636945 | GPCPD1 | T/C | -.20 | 1.4x10<sup>-6</sup> | -.10 | 2.5x10<sup>-2</sup> | -.03 | 4.2x10<sup>-7</sup> |
| **DLCO/VA\*** | rs17280293 | 6:142688969 | GPR126 | A/G | -.06 | 3.0x10<sup>-3</sup> | -.08 | **6.7x10<sup>-9</sup>** | -.01 | **1.4x10<sup>-10</sup>** |
|  | rs918606 | 5:61926379 | IPO11 | A/G | -.02 | 2.0x10<sup>-3</sup> | -.02 | 5.8x10<sup>-6</sup> | -.01 | 6.0x10<sup>-8</sup> |
|  | rs75834976 | 4:5231710 | STK32B | A/C | -.04 | 2.4x10<sup>-3</sup> | -.04 | 5.5x10<sup>-5</sup> | -.01 | 6.0x10<sup>-7</sup> |
|  | rs56315120 | 1:165168869 | LMX1A | A/G | -.02 | 0.24 | -.06 | 1.5x10<sup>-7</sup> | -.01 | 7.8x10<sup>-7</sup> |
| **DLCO/VA†** | rs17280293 | 6:142688969 | GPR126 | A/G | -.06 | 4.3x10<sup>-3</sup> | -.07 | **2.3x10<sup>-9</sup>** | -.01 | **7.9x10<sup>-11</sup>** |
|  | rs918606 | 5:61926379 | IPO11 | A/G | -.02 | 1.2x10<sup>-3</sup> | -.02 | 1.3x10<sup>-5</sup> | -.004 | 7.5x10<sup>-8</sup> |

A1: The first allele; A2: the second allele; B: the effect estimate which are additive effects for each copy of A1; Chr:Pos: Chromosome and position; DLCO: Diffusing capacity of carbon monoxide; DLCO/VA: Diffusing capacity of carbon monoxide by alveolar volume; FHS: the Framingham Heart Study; RS: the Rotterdam Study (Meta-analysis RSI, RSII and RSIII); SNP: single nucleotide polymorphism
\*Model 1: Adjusted for age, sex and principal components
†Model 2: Adjusted for age, sex, weight, height, smoking and principal components
‡The Gene name is a label of the region using the closest gene but does not necessarily pinpoint the responsible gene
Bold indicates statistically significant.

Analyses were adjusted for age, sex and PC in model 1. In model 2 analyses were adjusted for variables in model 1, in addition to weight, height, current and past smoking.

**Figure 2** represents the Manhattan-plots of DLCO GWAS at the meta-analysis level. For both DLCO analyses (models 1 and 2), no variant reached genome wide significance threshold. In model 2, two variants at 10q22.1 (rs1665630, gene: *CDH23*, MAF: 0.44, P-value= 2.8 × 10^−7^) and at 20p12.3 (rs2423124, close to gene: *GPCPD1*, MAF:0.19, P-value=4.2 × 10^−7^) showed a suggestive association with DLCO.

**Figure. 2:**
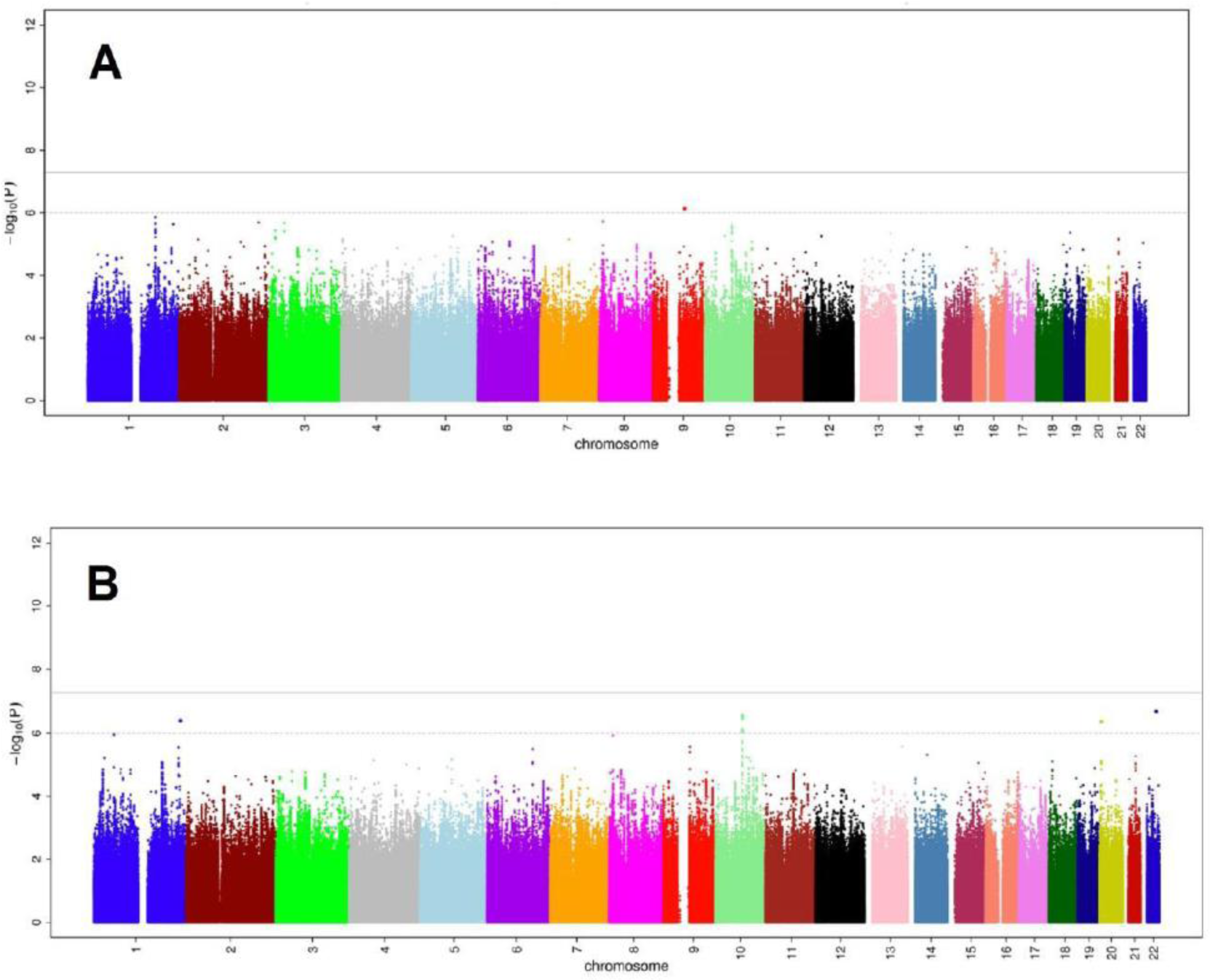
Common genetic variants associated with diffusing capacity of the lung for carbon monoxide (DLCO). A: Manhattan-plot of the association between common genetic variants and DLCO, adjusted for age, sex and principal components of genetic relatedness. B: Manhattan-plot of the association between common genetics variants and DLCO, adjusted for age, sex, weight, height, smoking and principal components of genetic relatedness.

**Figure 3** represents the Manhattan-plots of DLCO/VA GWAS at the meta-analysis level. Nineteen variants at the same locus at 6q24.1 (top: rs17280293, gene: *GPR126*; MAF: 0.03, P-value= 1.4 × 10^−10^), were significantly associated with DLCO/VA in model 1(see regional plot in **Figure 4).** Of these, six variants at the same locus at 6q24.1, reached the genome-wide significance threshold in model 2. Moreover, in both models, a variant at 5q12.1 (rs918606, gene: *IPO11*; MAF: 0.44; P-value model 1= 5.96 × 10^−8^, P-value-model 2= 7.49 × 10^−8^) was found to be suggestively associated with DLCO/VA. Sensitivity analyses by adjusting for haemoglobin blood concentrations or FEV_1_/FVC did not materially change the results of the DLCO/VA GWAS (see **Supplemental Results and Figure E2** in the Online Data Supplement).

**Figure. 3:**
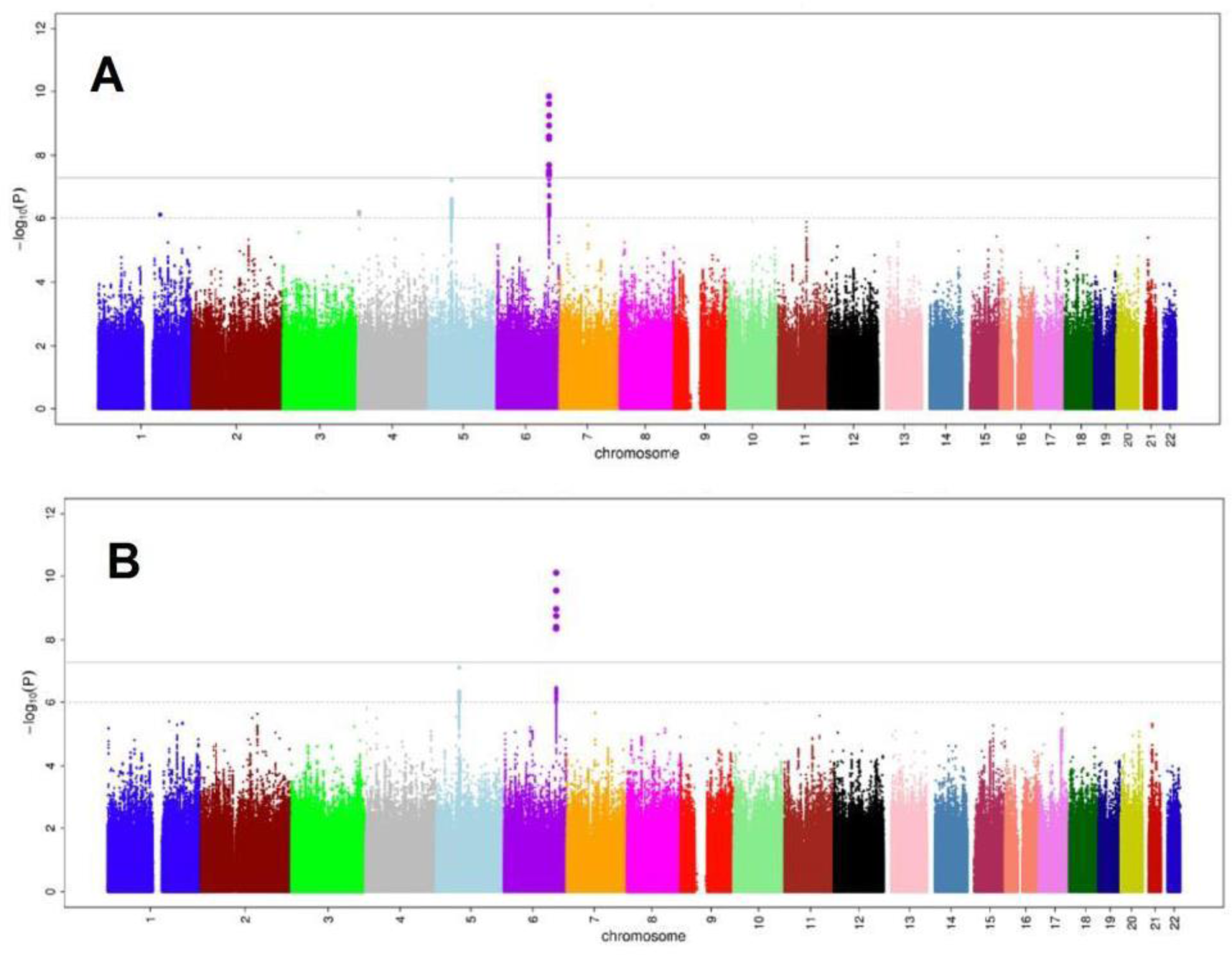
Common genetic variants associated with diffusing capacity of the lung per alveolar volume (DLCO/VA). A: Manhattan-plot of the association between common genetic variants and DLCO/VA, adjusted for age, sex and principal components of genetic relatedness. B: Manhattan-plot of the association between common genetics variants and DLCO/VA, adjusted for age, sex, weight, height, smoking and principal components of genetic relatedness.

**Figure. 4:**
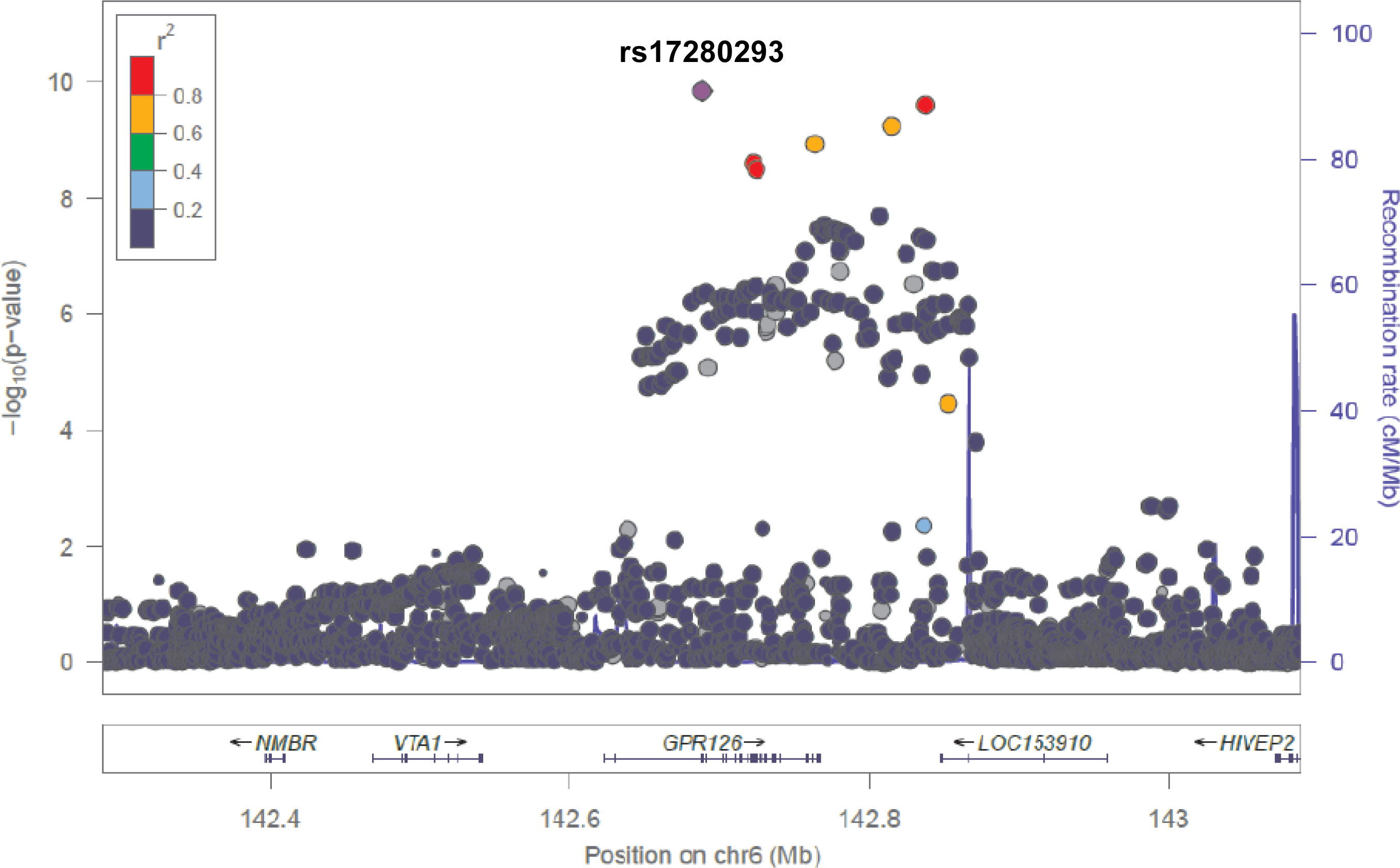
Regional association plot of the genome-wide significant locus in DLCO/VA GWAS.

Interestingly, a more in depth investigation of the *GPR126* region (**Figure 4**) revealed the presence of two missense variants; the lead SNP rs17280293 and rs11155242 (MAF 0.19, P-value=2.1 × 10^−06^). Those two SNPs showed to be in LD with each other, with r^2^=0.14 and D’=1.

### Follow-up analyses

In this section, the most important findings of the follow-up analyses will be summarized including genetic correlations and gene expression in lung tissue. Additional results on the genetic correlations, overlap with reported COPD and emphysema GWAS associations, posterior probability of causality, functional annotation and gene expression will be presented in the **Supplemental Results and Figures** in the Online Data Supplement.

### Genetic correlations

We examined the genetic correlation between DLCO/VA and DLCO using the age, sex, smoking status, weight, height and PC adjusted model. The genetic correlation was 59% (ρgenetic=0.59, P-value=0.04). We also examined the genetic correlation with FEV_1_/FVC and height (see **Supplemental Methods** and **Results** in Online Data Supplement).

### GPR126 expression

We extracted mRNA from lung resection specimens of 92 patients who underwent surgery for solitary pulmonary tumours or lung transplantation, including 44 patients without COPD and 48 patients with COPD (**Table 4**). The mRNA expression of *GPR126* was significantly lower in lung tissue of patients with decreased DLCO/VA compared with patients with normal DLCO/VA (**Figure 5A**) and in subjects with COPD (encompassing different categories of COPD severity according to the GOLD spirometric classification) compared to never smoking controls (**Figure 5B)**. The *GPR126* mRNA levels were significantly associated with DLCO/VA after adjustment for age and sex in model 1 (n=67 β=0.85 (95% CI 0.06-1.64)) and after additional adjustment for weight, height and smoking in model 2 n=66 (β=0.75 (95% CI 0.03-1.47)).

**Table 4:**
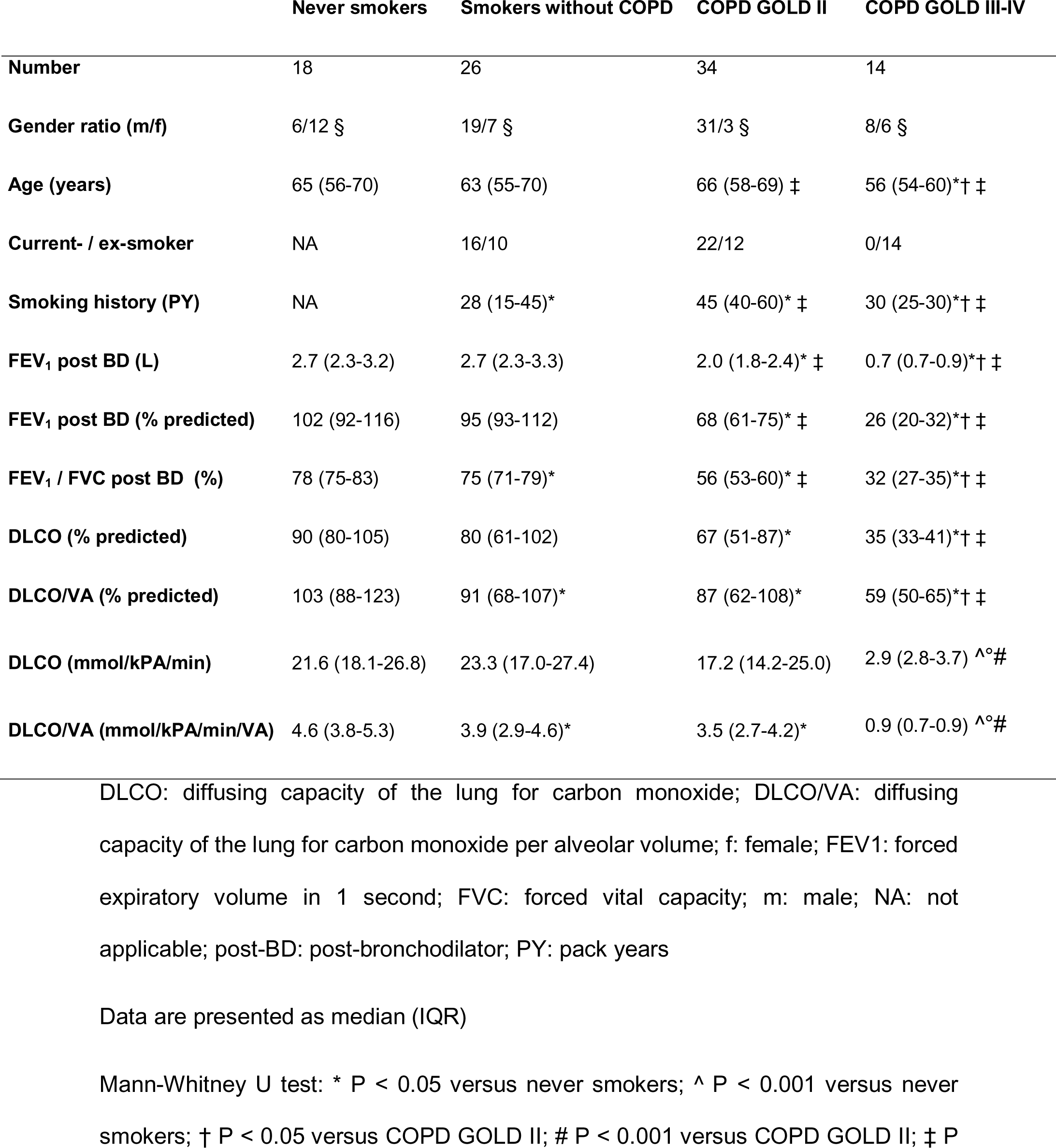
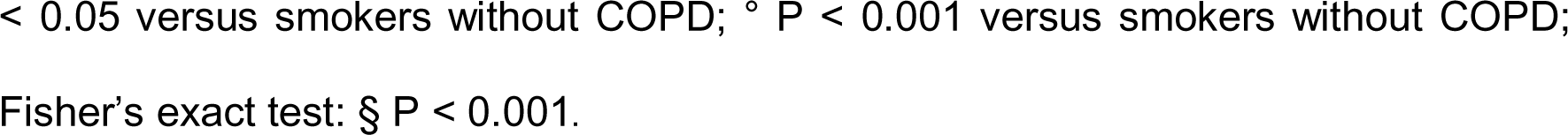
Characteristics of study individuals for lung mRNA analysis (by RT-PCR) (n=92)

|  | Never smokers | Smokers without COPD | COPD GOLD II | COPD GOLD III-IV |
| --- | --- | --- | --- | --- |
| **Number** | 18 | 26 | 34 | 14 |
| **Gender ratio (m/f)** | 6/12 § | 19/7 § | 31/3 § | 8/6 § |
| **Age (years)** | 65 (56-70) | 63 (55-70) | 66 (58-69) ‡ | 56 (54-60)*† ‡ |
| **Current- / ex-smoker** | NA | 16/10 | 22/12 | 0/14 |
| **Smoking history (PY)** | NA | 28 (15-45)* | 45 (40-60)* ‡ | 30 (25-30)*† ‡ |
| **FEV<sub>1</sub> post BD (L)** | 2.7 (2.3-3.2) | 2.7 (2.3-3.3) | 2.0 (1.8-2.4)* ‡ | 0.7 (0.7-0.9)*† ‡ |
| **FEV<sub>1</sub> post BD (% predicted)** | 102 (92-116) | 95 (93-112) | 68 (61-75)* ‡ | 26 (20-32)*† ‡ |
| **FEV<sub>1</sub> / FVC post BD (%)** | 78 (75-83) | 75 (71-79)* | 56 (53-60)* ‡ | 32 (27-35)*† ‡ |
| **DLCO (% predicted)** | 90 (80-105) | 80 (61-102) | 67 (51-87)* | 35 (33-41)*† ‡ |
| **DLCO/VA (% predicted)** | 103 (88-123) | 91 (68-107)* | 87 (62-108)* | 59 (50-65)*† ‡ |
| **DLCO (mmol/kPA/min)** | 21.6 (18.1-26.8) | 23.3 (17.0-27.4) | 17.2 (14.2-25.0) | 2.9 (2.8-3.7) ^°# |
| **DLCO/VA (mmol/kPA/min/VA)** | 4.6 (3.8-5.3) | 3.9 (2.9-4.6)* | 3.5 (2.7-4.2)* | 0.9 (0.7-0.9) ^°# |

DLCO: diffusing capacity of the lung for carbon monoxide; DLCO/VA: diffusing capacity of the lung for carbon monoxide per alveolar volume; f: female; FEV<sub>1</sub>: forced expiratory volume in 1 second; FVC: forced vital capacity; m: male; NA: not applicable; post-BD: post-bronchodilator; PY: pack years
Data are presented as median (IQR)
Mann-Whitney U test: \* P < 0.05 versus never smokers; ^ P < 0.001 versus never smokers; † P < 0.05 versus COPD GOLD II; # P < 0.001 versus COPD GOLD II; ‡ P
< 0.05 versus smokers without COPD; ° P < 0.001 versus smokers without COPD;
Fisher's exact test: § P < 0.001.

**Figure. 5:**
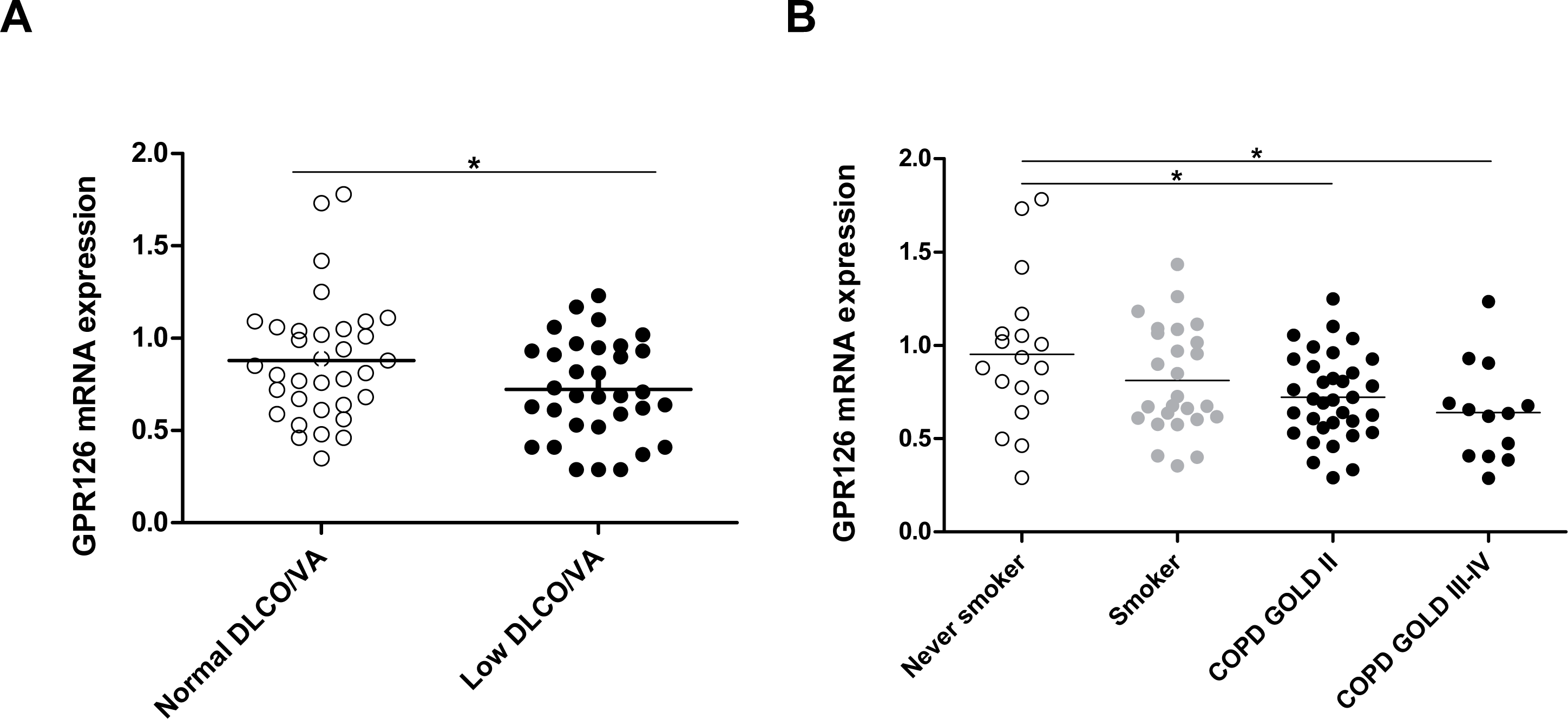
Pulmonary mRNA expression of *GPR126* in human subjects A. mRNA levels of *GPR126* in lung tissue of individuals with normal DLCO/VA (n=38) and low DLCO/VA (n=39). mRNA levels were corrected using a calculated normalization factor based on mRNA expression of three reference genes (GAPDH, SDHA, HPRT-1). B. mRNA levels of *GPR126* in lung tissue of never smokers (N=18), smokers without airflow limitation (N=26), patients with COPD GOLD II (N=34) and patients with COPD GOLD III-IV (N=14), as measured by quantitative RT-PCR. For statistical analysis, Kruskal-Wallis followed by Mann-Whitney U test was used for COPD (*P<0.05 and **P<0.01) and an independent sample t-test was used for DLCO/VA after rank transformation (* p<0.05).

## Discussion

This is the first study that has investigated the heritability of, and genome-wide association with, diffusing capacity of the lung using population-based cohort studies. We found a considerable proportion of variance in diffusing capacity of the lung explained by genetics. We also identified one locus on chromosome 6, encompassing the *GPR126* gene, that is associated with DLCO/VA and its lead variant showed to have a high posterior probability of causality compared to other SNPs in the same region. Finally, we were able to link the pulmonary expression of *GPR126* directly to COPD and to low DLCO/VA (compatible with emphysema in this general population). Here, we demonstrated a differential mRNA expression of *GPR126* in lung tissue of COPD patients and patients with decreased DLCO/VA.

### Heritability and genetic overlap

Studies on heritability of DLCO in the general population and unrelated individuals are lacking, and so far, DLCO heritability has been studied only in twins (15, 16), with a highest reported estimate of 44%. In our study, we estimated the REML based heritability of DLCO using the GCTA tool in unrelated individuals of the Rotterdam Study (17), and observed an age- and sex-adjusted heritability of DLCO and DLCO/VA of 23% and 24%, respectively. We also investigated heritability based on known familial relationships in the Framingham Heart Study. Here we found an age and sex-adjusted heritability of DLCO and DLCO/VA of 49% and 45%, respectively. The latter heritability estimates among familial related individuals are in line with the heritability estimates in twin studies and highlight the robustness of our data.

Importantly, our study is the first to investigate the lower bound of heritability of DLCO that would be estimated by family and twin studies (18). The advantage of estimating heritability in unrelated individuals using GCTA in addition to the approach based upon family and twin studies is, that GCTA calculates the proportion of heritability that covers the additive effects of commons SNPs only, and does not suffer from bias due to epistatic interactions or shared environment. The latter effects might indeed be present in family and twin studies, leading to an overestimation of the heritability (18-20).

Despite their similar estimates of heritability, DLCO and DLCO/VA showed to have different genetic determinants due to a genetic overlap between the two traits of 59%, explaining why we could not observe the same lead association in the two analyses.

### Variation in GPR126

The meta-analysis of genetics variants of DLCO/VA yielded one genome-wide significant association, along with a number of suggestive associations that did not reach genome-wide significance. The lead variant (rs17280293) in this study is a missense SNP in *GPR126*, with a MAF of 0.03 which is comparable to that in public datasets (0.03 ExAC, 0.02 TOPMED and 0.03 in 1000 genomes). Mutation in this SNP causes an amino acid change (S123G), which is predicted to have a deleterious effect as indicated by both SIFT (21) and Polyphen2 (22). It is therefore likely that this SNP is functional in *GPR126*. In this study, we showed that this variant has a high posterior probability of causality compared to other SNPs in the same region and that this SNP is associated with different regulatory chromatin marks, promotor histone marks, and enhancer histone marks in different tissue cell lines including foetal lung fibroblast cell lines and lung carcinoma cell lines. In addition, rs17280293 always co-occurs with another functional SNP in the region (rs11155242, D’=1), which is an eQTL for *GPR126* in human lung tissue.

Previous studies have also shown that variation at *GPR126* is associated with spirometric measures of lung function (5) (7). Soler Artigas and colleagues observed a strong association between spirometry, particularly FEV_1_/FVC, and another SNP rs148274477, which is in strong LD with rs17280293. However, since airflow limitation (i.e. a low FEV1/FVC ratio) might be correlated with low diffusing capacity due to loss of elastic recoil in subjects with emphysema, we assessed the possibility that the observed association between rs17280293 and DLCO/VA might be driven by FEV_1_/FVC. However, this sensitivity analysis indicated an independent association between rs17280293 and DLCO/VA because additional adjustment for FEV_1_/FVC did not affect the estimate and no genetic overlap could be proven between DLCO/VA and FEV_1_/FVC. Other studies have associated genetic variation in *GPR126* with height. In our study, adjustment for height did not affect the association between DLCO/VA and rs17280293, suggesting that the lead association in our GWAS is independent of height. In addition, genetic overlap disappeared after additional adjustment for height in the model, indicating no residual confounding by height in our analyses.

Furthermore, Eichstaedt and colleagues recently used whole genome sequence data from 19 Argentinean highlanders compared to 16 native American lowlanders and showed that rs17280293 might contribute to the physiological adaptations to hypobaric hypoxia (23).

### Gene function and expression

The *GPR126* gene belongs to the G-protein coupled receptor (GPCR) super family, the largest known receptor family in the human genome. It has been previously shown to be essential in angiogenesis (24). *GPR126*, a relatively new adhesion GPCR, has been shown to promote vascular endothelial growth factor (VEGF) signaling, by modulating the expression of endothelial growth factor receptor 2 (VEGFR2). Since *GPR126* is involved in angiogenesis, which is critical for the development of pulmonary capillary beds during fetal life, deletion of *GPR126* leads to mid-gestation embryonic lethality due to failure in cardiovascular development. GWA studies of spirometric measures of airflow limitation (FEV_1_/FVC ratio) have indicated several genes and pathways involved in branching morphogenesis and lung development, implicating an early life origin of complex adult respiratory diseases such as COPD. Intriguingly, this GWA study of diffusing capacity of the lung (DLCO and DLCO/VA) also indicates a gene (*GPR126*) which is implicated in cardiopulmonary development during fetal life.

The modulating effect of *GPR126* on *VEGFR2* expression was shown to be mediated through the transcriptional activation of *STAT5* and *GATA2* (24). Interestingly *GATA2* was recently linked to pulmonary alveolar proteinosis (25), a rare lung disease, characterized by an abnormal accumulation of pulmonary surfactant in the alveoli, leading to an altered gas exchange.

Moreover, knock down of *GPR126* in the mouse retina was shown to result in the suppression of hypoxia-induced angiogenesis (24). This information is interesting in two ways: first it links *GPR126* to hypoxia, which is very much related to gas-exchange. Second, processes in the retina might provide a unique insight into lung microvasculature, since vascular changes in both the retina and the alveoli reflect very much the same process, i.e. micro-angiopathy.

Although there is a good body of evidence that *GPR126* is important in lung development and micro-angiopathy, mRNA expression of *GPR126* has not been studied in lung diseases such as COPD and decreased diffusing capacity. Therefore we performed an expression analysis of *GPR126* in human lung tissue and demonstrated that mRNA expression of *GPR126* is decreased significantly in patients with COPD and individuals with a decreased DLCO/VA.

### Strengths and limitations

We conducted our analyses using data from two population based studies; the Rotterdam Study and the Framingham Heart Study. The strength of these studies is the population-based setting including data from smokers and non-smokers, and the standardized prospective data collection. We are not aware of other population-based cohort studies that have DLCO data in genotyped individuals available. Therefore, replication in other population-based cohorts was not possible. Yet, the results of the independent analyses in the Rotterdam study and the Framingham Heart Study show, that rs17280293 already reaches genome-wide significance in the Framingham Heart Study and replicates in the Rotterdam Study. Finally, a gene expression analysis on lung tissue was performed in our lab in very well-defined patient groups.

This study has also some limitations. First, for the measurements of diffusing capacity, the single-breath technique was used. This technique is known to underestimate measurements of alveolar volume in individuals with obstructive disease or air trapping, since diffusing capacity cannot be measured in poorly ventilated areas of the lung. It is also known that the underestimation of VA will be greater in more severe COPD and less in milder COPD. However, in our population-based cohorts, there are few individuals with severe COPD; therefore, reducing the impact of the underestimation of VA in our study. Secondly, haemoglobin corrected DLCO measures were only available in the Rotterdam Study. However, the performed sensitivity analysis with or without correction for haemoglobin did not materially change the results within the Rotterdam Study. The high D’ between rs17280293 and rs11155242 might suggest linked variant occurrence. However, the high D’ between those variants –estimated using data from the 1000 genomes reference panel-could result from the inflated estimation of D’ due to the low frequency of the SNPs. For this, it would be helpful to estimate the D’ in a bigger reference panel such as the haplotype reference consortium whenever this information would be available in the future.

In conclusion, DLCO and DLCO/VA are heritable traits with a considerable proportion of variance in diffusing capacity of the lung explained by genetics. We identified a functional variant in *GPR126*, a gene which is involved in gas exchange and hypoxia and differentially expressed in lung tissue of patients with COPD and subjects with decreased diffusing capacity. Therefore, experimental studies are needed to investigate the pathophysiological mechanisms and their therapeutic implications.

### URLs

METAL,http://www.sph.umich.edu/csg/abecasis/metal/.

GCTA, http://cnsgenomics.com/software/gcta/

Locuszoom plots, http://locuszoom.org/

Genetic correlation-LDscore, https://github.com/bulik/ldsc

Haploreg, http://archive.broadinstitute.org/mammals/haploreg/haploreg.php

GTEx portal, http://www.gtexportal.org/home/

GTEx portal eQTL data, lung tissue set obtained from this location:

javascript:portalClient.browseDatasets.downloadFile(’Lung.allpairs.txt.gz’,’gtex_analy

sis_v7/single_tissue_eqtl_data/all_snp_gene_associations/Lung.allpairs.txt.gz’)

## Data availability

Data can be obtained upon request. Requests should be directed towards the management team of the Rotterdam Study, which has a protocol for approving data requests. Because of restrictions based on privacy regulations and informed consent of the participants, data cannot be made freely available in a public repository.

## Acknowledgement

The authors are grateful to the study participants, the staff from the Rotterdam Study and the participating general practitioners and pharmacists.

The generation and management of GWAS genotype data for the Rotterdam Study (RS I, RS II, RS III) was executed by the Human Genotyping Facility of the Genetic Laboratory of the Department of Internal Medicine, Erasmus MC, Rotterdam, The Netherlands. We thank Pascal Arp, Mila Jhamai, Marijn Verkerk, Lizbeth Herrera and Marjolein Peters, MSc, and Carolina Medina-Gomez, MSc, for their help in creating the GWAS database, and Karol Estrada, PhD, Yurii Aulchenko, PhD, and Carolina Medina-Gomez, MSc, for the creation and analysis of imputed data.

## Supporting information

### Supplemental methods

Rotterdam study
Framingham Heart Study
Lung function
Genetics
Heritability
Genetic correlations
Overlap with reported COPD and emphysema GWAS associations
FINEMAP
GPR126 mRNA expression in lung tissue of patients with or without COPD

Human study populations
Definitions
RNA extraction and real-time PCR-analysis
Statistical analysis

### Supplemental results

Haemoglobin-adjusted analysis
FEV1/FVC adjusted analysis
Follow-up analyses

Genetic correlations
Overlap with reported COPD and emphysema GWAS associations
FINEMAP
Functional annotation
GPR126 expression

### Supplementary tables and figures

**Figure E1** Quantile-quantile plot at meta-analysis level

**Table E1** Main results of the meta-analysis in the Rotterdam Study (RS)

**Table E2** Main results of the GWAS in the Framingham Heart Study (FHS)

**Figure E2** Double Manhattan-plot where results of the meta-analysis of DLCO/VA before and after adjustment with FEV_1_/FVC in model 2.

**Table E3** Overlap with reported COPD-associated variants and those associated with DLCO in Model 1

**Table E4** Overlap with reported COPD-associated variants and those associated with DLCO in Model 2

**Table E5** Overlap with reported COPD-associated variants and those associated with DLCO/VA in Model 1

**Table E6** Overlap with reported COPD-associated variants and those associated with DLCO/VA in Model 2

**Table E7** Overlap with reported emphysema-associated variants and those associated with DLCO and DLCO/VA in the models 1 and 2

**Figure E3** Haploreg analysis of the main results of the meta-analysis

**Figure E4** GTEx output of *GPR126* expression in different tissues

**Figure E5** The genotypes of rs17280293 and rs11155242 in GTEx lung tissue database.

